# Classifying cells with Scasat - a tool to analyse single-cell ATAC-seq

**DOI:** 10.1101/227397

**Authors:** Syed Murtuza Baker, Connor Rogerson, Andrew Hayes, Andrew D. Sharrocks, Magnus Rattray

## Abstract

**Motivation:** The assay for transposase-accessible chromatin using sequencing (ATAC-seq) reveals the landscape and principles of DNA regulatory mechanisms by identifying the accessible genome of mammalian cells. When done at single-cell resolution, it provides an insight into the cell-to-cell variability that emerges from identical DNA sequences by identifying the variability in the genomic location of open chromatin sites in each of the cells. Processing of single-cell ATAC-seq requires a number of steps and a simple pipeline to processes and analyse single-cell ATAC-seq is not yet available.

**Results:** This paper presents ScAsAT (single-cell ATAC-seq analysis tool), a complete pipeline to process scATAC-seq data with simple steps. The pipeline is developed in a Jupyter notebook environment that holds the executable code along with the necessary description and results. For the initial sequence processing steps, the pipeline uses a number of well-known tools which it executes from a python environment for each of the fastq files. While functions for the data analysis part are mostly written in R, it is robust, flexible, interactive and easy to extend. The pipeline was applied to a single-cell ATAC-seq dataset in order to identify different cell-types from a complex cell mixture. The results from Scasat showed that open chromatin location corresponding to potential regulatory elements can account for cellular heterogeneity and can identify regulatory regions that separates cells from a complex population.

**Availability:** The jupyter notebook with the complete pipeline applied to the dataset published with this paper are publicly available on the Github (https://github.com/ManchesterBioinference/Scasat). An additional notebook is also provided for analysis of a publicly available dataset. The fastq files are submitted at ArrayExpress database at EMBL-EBI (www.ebi.ac.uk/arrayexpress) under accession number E-MTAB-6116.

**Supplementary information:** Supplementary data are available at *bioRxiv* online.

## 1 Introduction

Single-cell epigenomics studies the mechanisms that determine the state of each individual cell of a multicellular organism (Schwartzman and Tanay, 2015). Assay for transposase-accessible chromatin (ATAC-seq) can uncover the accessible region of a genome by identifying open chromatin regions using a hyperactive prokaryotic Tn5-transposase (Buenrostro *et al*., 2013, 2015b). In order to be active in transcriptional regulation, regulatory elements within chromatin have to be accessible to DNA-binding proteins (Tsompana and Buck, 2014). Thus chromatin accessibility is generally associated with active regulatory elements that drive gene expression and hence ultimately dictates cellular identity. As the Tn5 transposase only binds to DNA that is relatively free from nucleosomes and other proteins, it can reveal these open locations of chromatin (Buenrostro *et al*., 2013).

Epigenomics studies based on bulk cell populations have provided major achievements in making comprehensive maps of the epigenetic makeup of different cell and tissue types (Farh *et al*., 2015; Gjoneska *et al*., 2015). However such approaches perform poorly with rare cell types and with tissues that are hard to separate yet consist of a mixed population (Schwartzman and Tanay, 2015). Also, as seemingly homogeneous populations of cells show marked variability in their epigenetic, transcription and phenotypic profiles, an average profile from a Bulk population would mask this heterogeneity (Shalek *et al*., 2013). Single-cell epigenomics has the potential to alleviate these limitations leading to a more refined analysis of the regulatory mechanisms found in multicellular eukaryotes (Macaulay and Voet, 2014).

Recently, the ATAC-seq protocol was modified to apply with single-cell resolution (Buenrostro *et al*., 2015b; Cusanovich *et al*., 2015). Buenrostro *et al*. (2015b) used a microfluidic approach to isolate the cells whereas Cusanovich *et al*. (2015) avoided physical isolation of cells by using a combinatorial indexing strategy. However, neither of the studies developed a clear bioinformatics pipeline for the processing of the data and its downstream analysis. Schep *et al*. (2017) recently introduced *chromVar* to analyse scATAC-seq data. Rather than considering the full list of chromosomal locations, analysis in chromVar depends on the loss or gain of chromatin accessibility on a set of genomic features which could be either motif positions or genomic annotations. SCRAT (Ji, 2017) also uses a number of predefined features like Motif, Encode cluster, Gene and Gene sets to cluster cells into different sub-populations. This limits its application to chromosomal locations that represent an annotated feature. Zamanighomi *et al*. (2017) introduced scABC that removes the requirement of predefined accessible chromatin sites and depends on the patterns of read counts mapped to open genomic regions for the unsupervised clustering of the cells (Zamanighomi *et al*., 2017). However, in scATAC-seq, chromosome accessibility is essentially a binary phenomenon due to the uniqueness of a chromosomal location and statistical approaches that are more appropriate for binary data should be used. Although in principle *Cicero* (Pliner *et al*., 2017) can work with a binary matrix directly, the authors aggregated the binary profile of the cells that are highly similar based on clustering or psuedotemporal ordering to generate a count matrix. This count matrix was then used to connect the regulatory elements to their target genes. Furthermore, most of these tools do not have a processing pipeline and are not flexible enough to modify the functionality according to user requirements. A bioinformatics pipeline that has the processing steps for scATAC-seq data, works directly on binary data and is flexible enough to easily incorporate user defined functionality is not yet available.

In this paper we introduce a bioinformatics pipeline to conduct the following tasks:

- Processing: Adapter trimming, quality control, mapping, removing blacklisted reads, removing PCR dupicates and calling peaks.
- Peak accessibility for each cell: We first merge all the single-cell BAM files to create a reference set of peaks. An accessibility matrix is then generated using this reference set with the accessibility information for these peaks in each of the cells.
- Quality Control: Lower quality cells and peaks are removed. Also, peaks that would cause stronger batch effects are removed.
- Downstream analysis: Clustering the cells, deconvoluting cell types from a mixed cell population or identifying differentially accessible peaks between two groups of cells.

Figure (1) describes the complete work-flow of Scasat. It starts with the processing steps followed by the downstream analysis of the single-cell ATAC-seq. In the following we provide details of the workflow and demonstrate its utility by applying this to deconvoluting a mixture of three oesophageal tissue (ie one is “normal” and two are “tumours”) derived cell lines.

**Figure 1:**
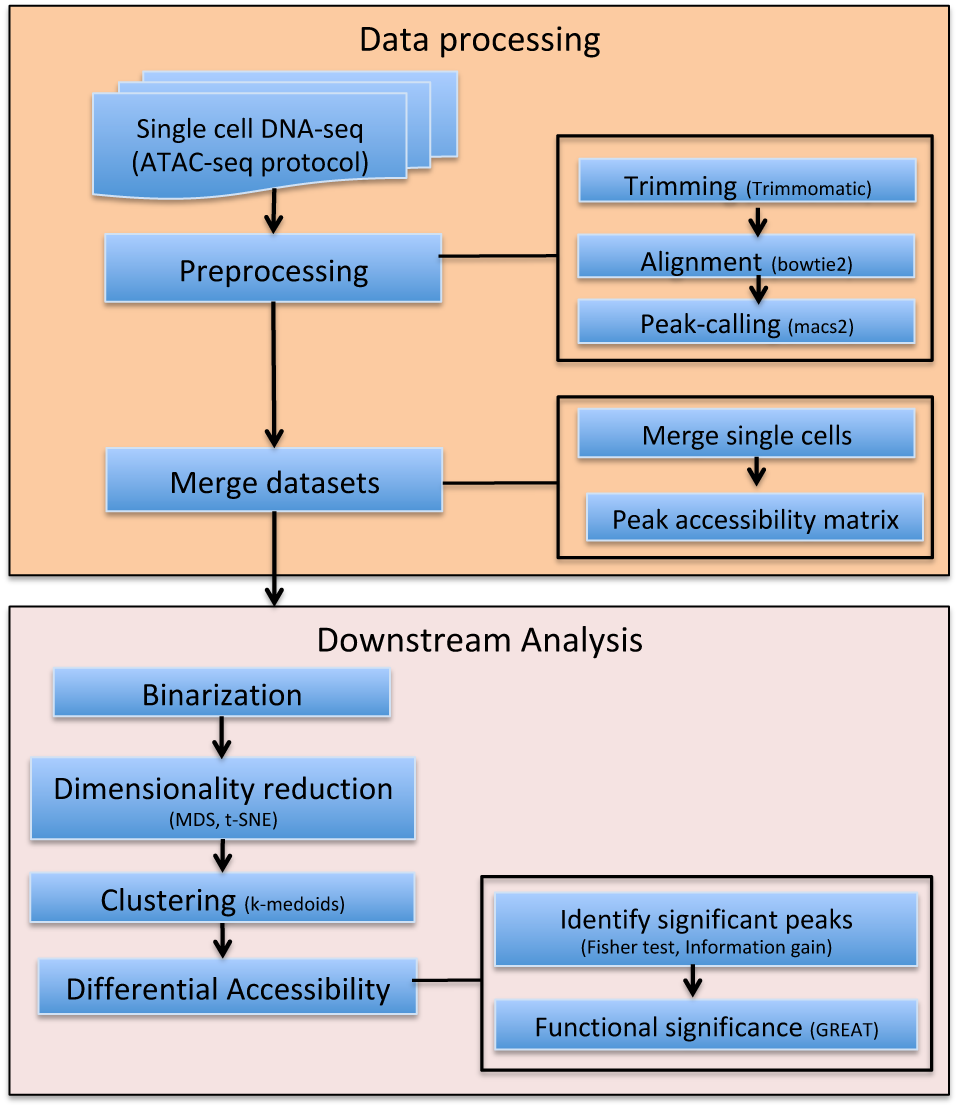
Workflow of the Scasat. It. shows the preprocessing as well as the downstream analysis available at Scasat

## 2 Method

Here we describe, Scasat, **S**ingle-cell **A**TAC-seq **A**nalysis **T**ools, processing and analysing single-cell ATAC-seq data. The Scasat workflow typically consists of four steps, 1. Data processing; 2. Feature extraction; 3. Heterogeneity analysis of the cells; 4. Differential accessibility analysis of the peaks between two cluster of cells. To make it convenient for the user, we have introduced two notebooks for this analysis. The first notebook does the processing of the data and the second notebook does the downstream statistical analysis. Below we discuss the workflow of both these notebooks.

### 2.1 Notebook environment

The complete pipeline was developed using the jupyter notebook environment (Pérez and Granger, 2007). The jupyter notebook is a web-based notebook that can execute code, produce figures and put all the necessary explanations in the same place. As all the function definitions of the tool are open to the user, it can be easily extended to integrate new tools or approaches. Although the current version of the pipeline enables only the serial processing of the cells at the processing step, it can be easily extended for parallel processing. The notebook makes it easy to document, understand and share the code with non-technical users (Shen, 2014). Scasat uses many of the most widely used software tools at the processing step. The parameters and paths of these tools are set in the python environment. For the downstream analysis Scasat uses the R programming languages for statistical computing and graphical visualization of the results. The use of R magic cells in the notebook variables makes the pipeline more robust and allows both programming languages to be used in the same workflow.

### 2.2 Sequence data processing

The processing step starts with first configuring the folders and setting the paths of the software. The user configures the *inputFolder* to the foldername where all the *fastq* files are. The *outputFolder* is configured to store all the processed files. Experiments using sequencing applications (ATAC-seq, Chip-seq) generate artificial high signals in some genomic regions due to inherent properties of some elements. In this pipeline we removed these regions from our alignment files using a list of comprehensive empirical blacklisted regions identified by the ENCODE and modENCODE consortia (Encode, 2012). The location of the reference genome is set through the parameter *ref_genome*. This folder contains the index file for the *bowtie2* aligner. A brief description of the tools that we have used in this processing notebook are given below

- Trimmomatic v0.36 (Bolger *et al*., 2014) is used to trim the illumina adapters as well as to remove the lower quality reads.
- Bowtie v2.2.3 (Langmead *et al*., 2009) is used to map paired end reads. We used the parameter −*X 2000* to allow fragments of up to 2kb to align. We set the parameter –dovetail to consider dovetail fragments as concordant. The user can modify these parameters depending on experimental design.
- Samtools (Li *et al*., 2009) is used to filter out the bad quality mapping. Reads with a mapping quality ¿ q30 are only retained. Samtools is also used to sort, index and to generate the log of mapping quality.
- Bedtools intersect (Quinlan and Hall, 2010) is used to find the overlapping reads with the blacklisted regions and then remove these regions from BAM file.
- Picards (Picard, 2017) MarkDuplicate is used to mark and remove the duplicates from the alignment.
- MACS2 (Zhang *et al*., 2008) is used with the parameters –nomodel, –nolambda, –keep-dup all –call-summits to call the peaks associated with ATAC-seq. During the callpeak we set the p-value to 0.0001. This is due to the fact that otherwise MACS2 will not call the peaks having a single read mapped to it as it would consider those reads to be background noise.
- Bdg2bw is used to generate the Bigwig files for the UCSC genome browser visualization.
- QC: A final quality control is performed based on the library size of the BAM file. We filter out the cells for which the library size estimated by Picard tool is less than a user-defined threshold. The default value of the LIBRARY_SIZE_THRESHOLD is set to 10000. We consider any cell having a library size lower than this threshold to not be a valid cell as those reads may come from debris free material or from dead cells.

### 2.3 Downstream analysis

Single-cell ATAC-seq is essentially binary in nature. A specific location in a chromosome for a specific cell can either be open or closed. In contrast to bulk data where more reads aligning to a specific location of a chromosome would indicate more cells in the population having open chromatin at that location, in single-cells it could only be due to the multiple insertions in that region or possibly other alleles at that locus. As described by Buenrostro *et al*. (2015b) such reads are rare for single-cell ATAC-seq which is overwhelmly dominated by single reads for a specific location of a chromosome for each individual cell. The analysis pipeline in Scasat is constructed taking into consideration the binary nature of chromatin accessibility in single cells. We merge the single-cells only to get the list of reference peaks. Scasat then binarizes this peak information for each individual cell for downstream analysis.

#### Peak accessibility matrix

The analysis workflow of Scasat starts by merging all the single-cell BAM files and creating a single aggregated BAM file. Peaks are called using MACS2 on this aggregated BAM file and sorted based on q-value. Peaks in this list are the ones that are open in at least one single-cell. Using this list of peaks we generate the *peak accessibility matrix*. The rows of this matrix represent all the peaks from the reference set and the columns represent each single cell. The pipeline calculates the accessibility of the peaks for each individual cell where it has at least one overlapping read and encodes it as a binary value. For each individual cell, peaks that overlap with this list of accessible regions are given the value of 1 in the table. For all the other peaks it is 0. A graphical explanation of this process is given in FIG. S1

#### Bulk vs aggregate

If a Bulk measurement is available for the same cell-type or sample, then the pipeline can calculate *Number_of_peaks vs. precision* for the aggregated single-cell data against its population-based Bulk data. This demonstrates how closely the single-cell data recapitulates its Bulk counterpart. We define *peakA’s* list all the peaks in the population based on Bulk data and *peakB’s* list the peaks in aggregated single-cells sorted on q-values. *peakA* is considered to be the gold standard for this calculation. We start with the top 100 peaks in the sorted peak list of *peakB*. We assume that all these 100 peaks are positive peaks overlapping with the peaks in *peakA*. Now, all the peaks within this 100 list that actually overlap with the peaks in *peakA* are considered the True Positive and the ones that do not overlap are the False Positive ones. Now, we take all the remaining peaks in *peakB* (peaks that we get after removing the 100 previously selected ones) as the negative ones. If any of the peaks in this negative set overlaps with peaks in *peakA* then we denote them as False Negative. Otherwise, they are called as True negative. We then calculate the precision as

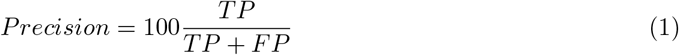

where, TP = True Positive, FP = False Positive, TN = True Negative, FN = False Negative. We then repeat the process by increasing *peakB* with 50 peaks each time while keeping *peakA* fixed. This gives us a *Recall vs Precision* plot show in FIG. S2.

#### Filtering Peaks

Peaks appearing in a small number of cells are less informative and are not always appropriate for downstream analysis. Similarly, some cells passing through library size filtering, might still have a very low number of peaks. In our downstream analysis we filter out these cells and peaks. If a cell has open peaks below a user defined threshold (default: 50 peaks or 0.2% of total peaks) in *peak accessibility matrix* we would drop that cell. Also peaks not observed across a user defined number of valid cells (default :8 cells) are not considered for further downstream analysis. Choice of this threshold depends on the number of cells in the experiment, nature of those cells and other biological as well as technical factors. Users need to carefully define these thresholds for the filtering based on their experimental design.

#### Calculate Jaccard distance

Once the accessibility matrix is generated, we are interested in a dissimilarity measure that quantifies the degree to which two cells vary in their peak accessibility. In our pipeline we use the Jaccard distance (Jaccard, 1901) as a dissimilarity measure. The Jaccard distance is the ratio between the number of peaks that are unique to a cell against all the peaks that are open in two cells.

#### Dimensionality reduction

Our *peak accessibility matrix* represents a very high-dimensional dataset of open chromatin regions for each single-cell. Dimensionality reduction for this high-dimensional dataset is essential for easy visualization and other downstream analysis. In our pipeline we applied the multidimensional scaling (MDS) and the t-distributed stochastic neighbour embedding (t-SNE). MDS provides a visual representation of similarity (or dissimilarity) between two objects. It takes as input the distances between any pair of objects and then minimizes a loss function called strain (Borg and Groenen, 2003) so that the between object distances are preserved as much as possible. t-SNE is a non-linear dimensionality reduction technique that maps multidimensional data to two or more dimensions for easy visualization. t-SNE converts the similarity between the data points to joint probabilities and tries to minimize the Kullback-Leibler divergence between the joint probabilities of high dimensional data and low-dimensional embedding of this data (van der Maaten and Hinton, 2008). To reduce the noise and computational speed it is recommended to use a lower dimensional representation of the data as input to t-SNE.

#### Clustering

In this pipeline we used the k-medoids algorithm to cluster the cells into different groups. The k-medoids algorithm breaks the dataset into different partitions and attempts to minimize the distance between points assigned in a cluster and a point designated as the center of that cluster. In our pipeline we used Jaccard distance as the dissimilarity matrix for this algorithm.

#### Differential accessibility (DA) analysis

One of the key features of interest in single-cell ATAC-seq analysis is to perform a statistical analysis to discover whether quantitative changes in accessibility of chromatin locations between two groups of cells are statistically significant. The pipeline implements two methods, Fisher exact test and Information gain to conduct the differential accessibility analysis between any two groups of cells.

#### Information gain

Based on the expected reduction in entropy (homogeneity measure), information gain measures the attribute that provides the best prediction of the target attribute where entropy is reduced. In differential accessibility analysis, *information gain* lists the peaks that best splits the cells into different groups. Defining Entropy as 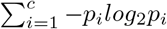, the Information gain is calculated as

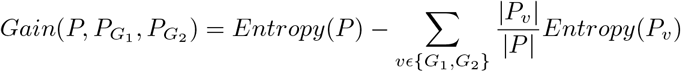

Here, *P* is the collection of all data, *P*_*G*_1__ represents the cells in *G*_1_ and *P*_*G*_2__ represents cells in *G*_2_. We calculate the information gain of each of the peaks given the cells are divided into two groups. The peaks are then sorted based on the informatin gain and the user can choose the cutoff value for selecting the DA peaks.

#### Fisher exact test

The Fisher exact test looks at a 2 × 2 contingency table that shows how different groups/conditions have produced different outcomes. Its *null hypothesis* states that the outcome is not affected by groups or conditions. We run this test on a peak-by-peak basis by organizing the open and closed (1’s and 0’s) for each peak in a 2 × 2 contingency table. The p-values are then corrected using Bonferroni correction for multiple comparisons (Bonferroni, 1936). Differentially accessible peaks with statistical significance are then selected based on a user-defined cutoff value (default: q-value ¡ 0.01).

## 3 Results

To demonstrate Scasat we generated scATAC-seq data from a mixture of three different cell types and the objective was to identify these three cell types from this mixture. We applied Scasat to characterize biologically relevant chromatin variability associated with each cell-type.

### Experimental design

To create the mixture we took two classic oesophageal adenocarcinoma (OAC) cell lines, OE19, OE33 and one non-neoplastic HET1A cell line. We mixed the three cell types in equal proportions to create a heterogeneous population. Two samples from this mixture were taken to make two technical replicates. ATAC-seq was then performed on those two replicates by loading on two separate C1 fluidigm chips using a 96 well plate integrated fluidic circuit (IFC) and sequenced on an Illumina NextSeq (Figure (2)). As ATAC-seq reports on the accessible regions of the chromatin which are considered to be active (Buenrostro *et al*., 2015a), these three cell lines are expected to have different accessibility at regulatory regions. The analysis attempted to disentangle these cells based on the presence of these active sites.

**Figure 2:**
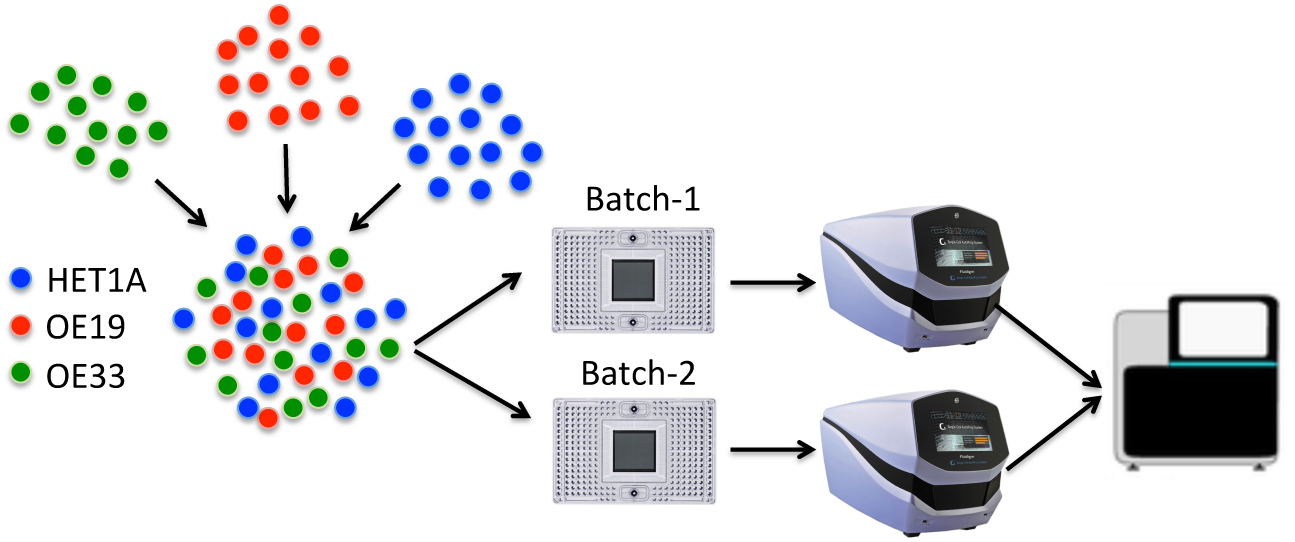
Experimental design of cell disentangle. Three cell types HET1A, 0E19 and OE33 are mixed into equal proportion. Samples are then taken from this mixture into two independent batches (Batch-1 and Batch-2). The cells are then sequenced using a NextSeq.

### Sample preparation

We plated 1 × 10^6^ of each cell type onto 10 *cm* plates and incubated for 24 hours at 37°*C*. Cells were detached using 0.1% trypsin. After detaching cells, each individual cell type was placed into a separate tube, centrifuged and re suspended in 1 *ml* of 1*X* phosphate-buffered saline (PBS). Each cell type suspension was quantified using a haemocytometer and this gives us a concentration in cells per *ml*. We then put the same number of each cell type into the same tube (to get the mixed population), which was then centrifuged again and re-suspended in 100 *μl* of 1*X PBS*. This was then submitted to the Genomic Technologies Core Facility. The C1 platform (Fluidigm) was used to capture single cells and generate sequencing libraries using the scATAC-seq protocol available from the Fluidigm ScriptHub (Buenrostro *et al*., 2017). Medium (10-17 micron) C1 for OpenApp Integrated Fluidic Circuits (IFCs) were used to capture and process the samples. The amplified DNA products were harvested, and then additional PCR performed to dual-index the harvested libraries using customized Nextera PCR primer barcodes (IDT Technologies) according to the ScriptHub protocol. The PCR products were then pooled to a total volume of 96*μl*, followed by two cycles of AMPure XP bead purification (Beckman Coulter, Inc.) according to the SMART-seq v4 protocol 032416 (Clontech Laboratories, Inc.) using C1 Dilution Reagent (Fluidigm) for the final elution. The pools were then quantified and validated using qPCR-based KAPA library quantification kit for Illumina (KAPA Biosystems) and TapeStation 4200 (Agilent Technologies). Library pools (1.8 pM final concentration) were then sequenced (75:75 bp, paired-end) on the NextSeq500 platform using the Mid Output v2 150 cycle kit (Illumina, Inc.) to generate .fastq sequence files.

### Data processing and analysis

Two batches were processed separately making it easier to keep track of the processing steps and to troubleshoot any problem that might arise due to batch effects. The parameters for the tools are explained in the tool description section. We trim the adapter sequences using Trimmomatic and used Bowtie2 to map reads to the genome. After removing the Blacklisted regions, we use *Picard’s Markduplicate*, to remove the duplicates. We then remove the chrY (as the three cells lines are a mixture of male and female) and chrM. The peak calling is done in two stages. In the first stage *macs2* is used to call peaks in each individual cell by setting the *p-value* parameter to 0.0001 to ensure that peaks associated with low mapping reads are also called. We then filter the cells that fail to cross the LIBRARY_SIZE_THRESHOLD set to 10000. After this QC, Batch-1 has 84 and Batch-2 had 89 cells for downstream analysis. In the second stage of peak calling, we aggregated all the BAM (both batches) files by calling *getAggregatedPeak()* module and use *macs2* to call peaks on this aggregated BAM file with qvalue of 0.2. This gave us a reference set of peaks. We then use the *mergePeaks()* module to merge the overlapping peaks in this reference set and sort them based on the q-value giving us a total of 236, 580 peaks as reference set. Finally we call *peakAccessibility()* module to calculate the accessibility of these reference peaks for each single-cell and generated the *peak accessibility matrix*.

### Peak selection

We used the *clean.count.peaks()* with the default parameters to remove the lower quality cells and peaks. When setting *min.cell.peaks.obs=10* (a valid peak has to be observed in at least 5% cells across both batches), a significant number of peaks were removed. These peaks are filtered out because we believe they do not contain enough information for reliable statistical inference. One caveat of this approach is that peaks associated with more abundant cell types are potentially kept and we might fail to detect rare sub-populations. In such cases the threshold for number of cells with a peak can be reduced.

### Dimensionality reduction

We now employed the *plotMDS()* module that subsequently calls the *getJaccardDist()* module to calculate Jaccard distances between the cells, applies multidimensional scaling on this Jaccard distance and plots the cells in a 2*D* scatter plot. Looking at the MDS plot we noticed a clear batch effect (Figure 3(A)).

**Figure 3:**
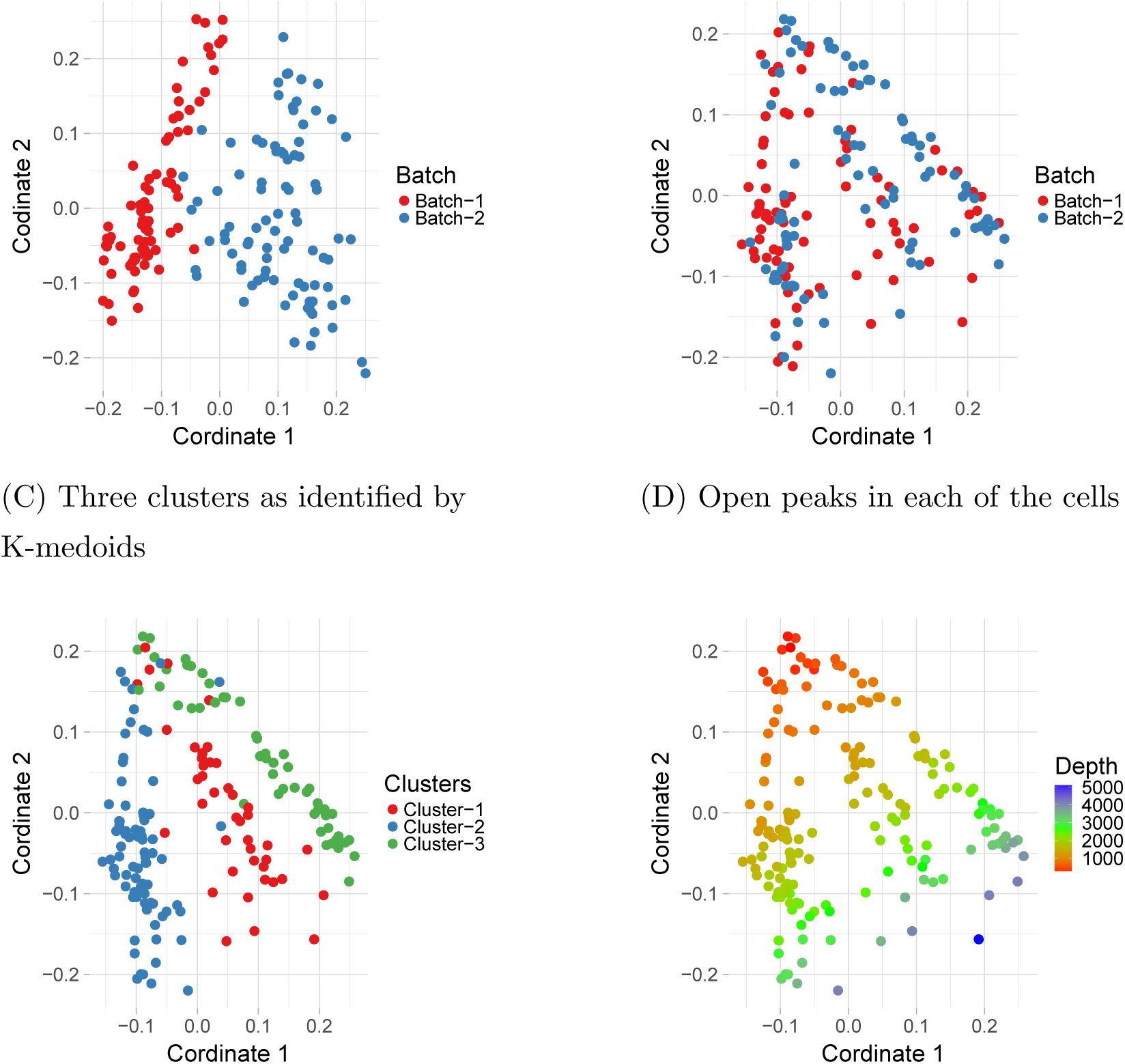
MDS plot with (A) and without (B) additional filtering to remove peaks where very few cells from a specific batch have open peaks. In (A) cells can be separated using only the batch information which is removed in (B) with additional filtering. (C) shows the three clusters identified by k-medoids clustering algorithm. (D) MDS plot with number of open peaks superimposed on them. Coordinate 2 is dominated by number of open sites per cell which is highly correlated with number of open peaks.

### Additional filtering

Applying conventional batch correction methods like linear regression as in Limma or other approaches outlined in SVA or Combat are not applicable here as all of them would convert the binary data into a real number. So we had to apply a different approach. Careful analysis of the data found that the peaks in Batch-1 had higher number of zeros associated with aggregated peaks, indicating that less information is attained from Batch-1 cells compared to Batch-2 (FIG S2). So we applied an additional filtering. In that filtering, if a peak is observed in less than three cells in any of the batches we discarded that peak. This further reduced the number of peaks to 17, 255 peaks. Applying MDS to these peaks the cells no longer show a strong batch effect (Figure 3(B)).

### Clustering

We used the *Partitioning Around Medoids (PAM)* algorithm which is the most common realisation of k-medoid clustering to cluster the cells. For this algorithm we chose *K* = 3 as we expected the cell-mixture to have three types of cells. While calculating the partitions we passed the Jaccard distance to the function instead of the actual dataset. The assignment of the clusters for each of the cells are superimposed onto our MDS plot which is shown in Figure 3(C). Figure 3(D) colors the cells based on the number of open peaks. Although the cells separate on Coordinate2 in Figure 3(D) based on the availability of the open peaks, they do not mask the actual clustering of the cells in Figure 3(C) as the cell-clustering is dominated with the distances in Coordinate1. Cluster1 has 42 cells, Cluster2 has 80 cells and Cluster3 has 51 cells.

### Identifying cell types

To relate the different groups to the input cell types, we compared it with the Bulk data of HET1A, OE19 and OE33 and also with single-cell datasets for OE19 (done in two batches, B1 and B2) and HET1A (Batch B1) by aggregating the single cells into its corresponding cell type. For this comparison, we merged the mapped reads for each cell in each of the clusters to create three aggregated BAM files, one for each of Cluster1, Cluster2 and Cluster3 identified in Figure 3(C). For the other two single-cell dataset we did the same. We now extend the column of *peak accessibility matrix* by calculating the peak accessibility for the three aggregated single cell data (OE19 B1, B2 and HET1A B1) and the Bulk datasets (HET1A, OE33, OE19) using the same 17, 255 peaks. Finally, we calculate the Pearson correlation coefficient for each of these dataset against each other which is shown in a correlation plot (Figure 4) and cluster them using hierarchical clustering. The correlation plot assigns each of the single cell clusters to their respective cell type based on the high correlation coefficient with the known cell types. This identifies Cluster1 as OE33 cell, Cluster2 as OE19 cell and Cluster3 as HET1A cell (Figure 4). If the Bulk data is not available or any other input cell types are not known, GO based analysis eg. GREAT (McLean *et al*., 2010) can be used to assign cells to a particular cell type.

**Figure 4:**
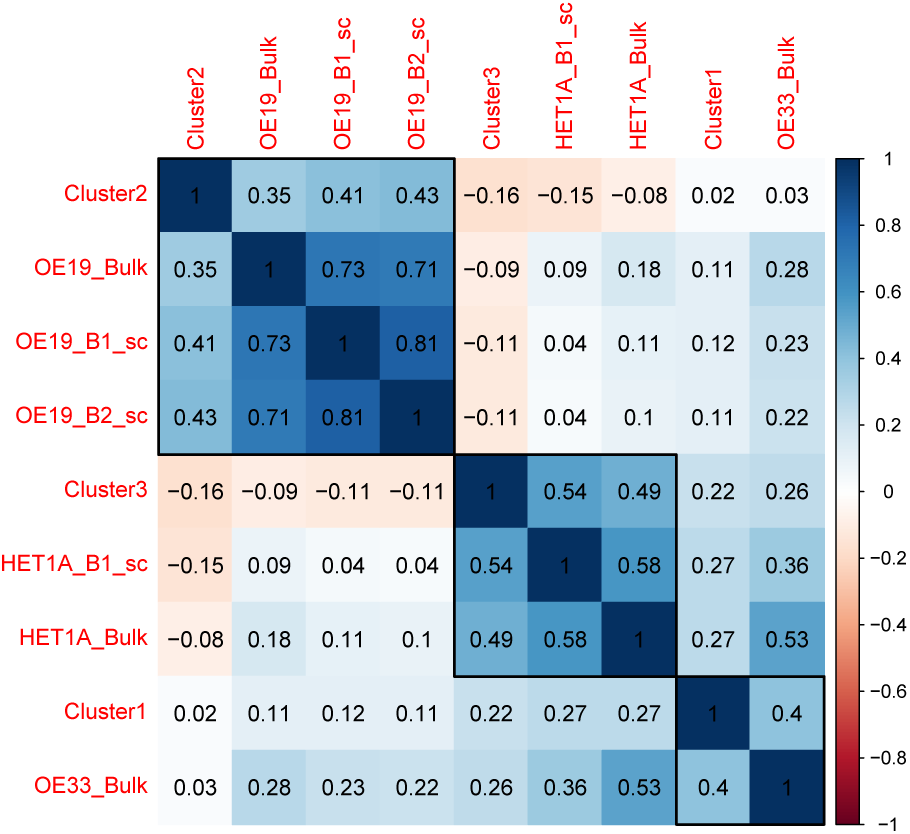
Correlation plot showing Pearson correlation for the three cluster of cells identified with k-medoid clustering algorithm, against the Bulk and two other single-cell data for which the cell types were known a priori. These known single-cell data are for HET1A done in one batch (B1) and OE19 done in two batches (B1 and B2). Single cells from these known cell types are aggreated and has label sc at the end in this plot. The bulk dataset have the label “_Bulk”. Here red indicates negative correlation and blue indicates positive correlation (scale bar on the right).

### Evaluating clustering solutions with golden truth

We used the Bulk data to identify the Ground truth on this data by calculating the correlation for each single-cell against the three Bulk data and assigning each cell to its corresponding type based on the largest correlation coefficient. These assignments are then projected onto MDS plot in FIG. S4. This Ground Truth shows very similar cell assignment as with our k-medoid algorithm. The adjusted rand index (ARI) that measures the similarity between two clusters gives a value of 0.75 for the cell assignment by k-medoid clusters and the ground truth where 1 signifies complete overlap between the two assignments.

### UCSC Genome tracks

We created a genome browser view of the aggregated single cells in the clusters and compared this to the data from Bulk ATAC-seq experiments performed on known cell populations. We ran our Differential Accessibility analysis with the module *getDiffAccessInformationGain()* which uses the entropy and information gain to identify the differentially accessible peaks between *Cluster1 vs Cluster2* and *Cluster2 vs Cluster3*. We annotated these peaks by finding the overlapping genes with a maximum distance of 1000 bases. Figure(5) shows the peak pattern for each group of cells and its known Bulk cell type. Two genes that have differentially accessible peaks between Cluster2 and Cluster3 are *GATA6* and *PPFIA3*. In Figure (5(A) we see that *GATA6* and *PPFIA3* are more open in Cluster2 and OE19 Bulk data confirming Cluster2 to be mostly comprised of OE19 cells. The peaks associated with *PPFIA3* gene in Figure (5(B)) also have very similar pattern in Cluster2 and OE19 cells. The activation of *GATA6* can sustain oncogenic lineage survival in esophageal adenocarcinoma (Lin *et al*., 2012) while *PPFIA3*, is a gene encoding a receptor that has been reported to show moderate cytoplasmic activity in colorectal cancers (Gene, 2017) which can develop into oesophageal metastasis (Thomasset *et al*., 2008). We find that *CAV1* is differentially accessible between Cluster1 and Cluster2 which is a tumour suppressor gene candidate (Trimmer *et al*., 2013) which have higher opening in Bulk OE33 cells and our Cluster1 cells (Figure(5C) confirming Cluster1 as mostly OE33 cells. Britton *et al*. (2017) reported an intragenic open chromatin location at the *MAS1* locus for HET1A (Britton *et al*., 2017) which we also observe in Figure (5D) for both our Cluster3 and HET1A cells, confirming the identify of Cluster3 cells mostly as HET1A cells.

**Figure 5:**
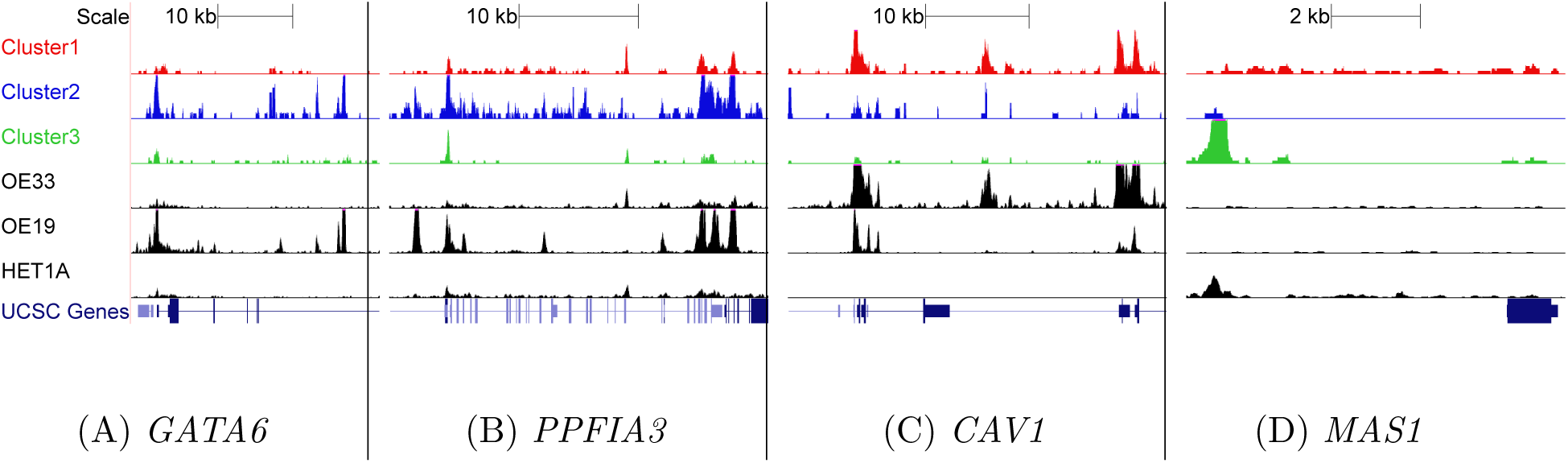
Confirmation of our assignments through visualizing peaks across different genome location from UCSC genome browser track. The clusters in three different colors (red, blue, green) corresponds to the inferred aggregated single-cell peak pattern. The bottom three peaks in Black shows the peaks for the Bulk data of OE33, OE19 and HET1A respectively. Peaks around the gene body of (A) GATA6, (B) PPFIA3, (C) CAV1, (D) MAS1 are shown in the figure. For these genes, peaks are more open in (A) & (B) OE19 cells, (C) OE33 (D) HET1A cells.

### GO based functionality

To confirm these cluster assignments further, we looked at the disease ontology associated with the cells in the clusters. We used the peaks that we identified during differential accessibility analysis of Cluster2 and Cluster3 and took the peaks that are differentially accessible with with more than two log2 fold change in Cluster2 compared to Clsuter3 and identified the Disease ontologies associated with these peaks through GREAT. GREAT first associate the peaks with potential target genes and then find out the disease ontologies associated with these open chromatin locations. The same process is repeated for Cluster3. We then identify the 15 topmost diseases associated with Cluster2 and maps the statistical significance for these disease in Cluster3. Although esophageal carcinoma did not make into the topmost 15 disease in Cluster2 (It came in 33^*th*^ position) we added it into this list of topmost genes. The −log10(Binom p-value) for these diseases are shown in FIG. S5. Esophageal carcinoma is picked up as one of the diseases associated with Cluster2 with very high confidence whereas the statistical significance is almost not present for Cluster3 cells. Several other significant disease associations are seen with adenocarcinomas and with general cancer. In almost all of these disease ontologies Cluster2 shows very high statistical significance whereas Cluster3 shows almost no association with very low p-value. This further confirms our accurate identification of HET1A cells and cancer derived subtypes supporting the conclusion of Corces *et al*. (2016) that scATAC-seq in addition to scRNA-seq can accurately identify cell types in complex cellular populations.

In summary, we showed that peak accessibility information from scATAC-seq can be used to deconvolute single-cells from a mixed cell population. We showed that an unsupervised clustering algorithm clusters cells according to their respective cell type. Similar methods could be applied to separate malignant cells from normal cell in a cancerous tissue and then investigate the malignant cells in detail.

## Conclusions

As single-cell ATAC-seq experiments are gaining momentum, we report Scasat, a pipeline to process and analyse single-cell ATAC-seq. Scasat offers two major utilities, the initial processing of scATAC-seq data and its downstream analysis. Scasat is implemented in jupyter notebooks making it simple, robust, scalable and easy to extend. Results from our data showed that an unsupervised clustering of the cells based on accessible chromatin regions can group cells into their corresponding cell type. This suggests that regulatory elements can define cell identity quite precisely. The successful implementation of this tool helped us to understand further the epigenetic mechanisms at the single-cell level and opens opportunities for easier and better analysis of single-cell ATAC-seq data.

## Acknowledgements

We thank Leo Zeef, Ian Donaldson (Bioinformatics core facility) for the technical discussions. We thank Edward Britton from Andy Sharrocks group for his discussion on ATAC-seq. We would also like to thank people in the Rattray lab for the critical evaluation and feedback on the tool.

## References

Bolger, A. M., Lohse, M., and Usadel, B. (2014). Trimmomatic: a flexible trimmer for illumina sequence data. Bioinformatics, 30(15), 2114–20.

Bonferroni, C. E. (1936). Teoria statistica delle classi e calcolo delle probabilita. Pubblicazioni del R Istituto Superiore di Scienze Economiche e Commerciali di Firenze, 8, 3–62.

Borg, I. and Groenen, P. (2003). Modern multidimensional scaling: Theory and applications. Journal of Educational Measurement, 40(3), 277–280.

Britton, E., Rogerson, C., Mehta, S., Li, Y., Li, X., the OCCAMS consortium, Fitzgerald, R. C., Ang, Y. S., and Sharrocks, A. D. (2017). Open chromatin profiling identifies ap1 as a transcriptional regulator in oesophageal adenocarcinoma. PLoS Genetics, 13(8), e1006879.

Buenrostro, J. D., Giresi, P. G., Zaba, L. C., Chang, H. Y., and Greenleaf, W. J. (2013). Transposition of native chromatin for fast and sensitive epigenomic profiling of open chromatin, dna-binding proteins and nucleosome position. Nat Methods, 10(12), 1213–8.

Buenrostro, J. D., Wu, B., Chang, H. Y., and Greenleaf, W. J. (2015a). Atac-seq: A method for assaying chromatin accessibility genome-wide. Curr Protoc Mol Biol, 109, 21 29 1–9.

Buenrostro, J. D., Wu, B., Litzenburger, U. M., Ruff, D., Gonzales, M. L., Snyder, M. P., Chang, H. Y., and Greenleaf, W. J. (2015b). Single-cell chromatin accessibility reveals principles of regulatory variation. Nature, 523(7561), 486–90.

Buenrostro, J. D., Giresi, P. G., Zaba, L. C., Chang, H. Y., and Greenleaf, W. J. (2017). Fluidigm c1-atac-seq scripthub.

Corces, M. R., Buenrostro, J. D., Wu, B., Greenside, P. G., Chan, S. M., Koenig, J. L., Snyder, M. P., Pritchard, J. K., Kundaje, A., Greenleaf, W. J., Majeti, R., and Chang, H. Y. (2016). Lineage-specific and single-cell chromatin accessibility charts human hematopoiesis and leukemia evolution. Nat Genet, 48(10), 1193–1203.

Cusanovich, D. A., Daza, R., Adey, A., Pliner, H. A., Christiansen, L., Gunderson, K. L., Steemers, F. J., Trapnell, C., and Shendure, J. (2015). Multiplex single cell profiling of chromatin accessibility by combinatorial cellular indexing. Science, 348(6237), 910–4.

Encode, P. C. (2012). An integrated encyclopedia of dna elements in the human genome. Nature, 489(7414), 57–74.

Farh, K. K., Marson, A., Zhu, J., Kleinewietfeld, M., Housley, W. J., Beik, S., Shoresh, N., Whitton, H., Ryan, R. J., Shishkin, A. A., Hatan, M., Carrasco-Alfonso, M. J., Mayer, D., Luckey, C. J., Patsopoulos, N. A., De Jager, P. L., Kuchroo, V. K., Epstein, C. B., Daly, M. J., Hafler, D. A., and Bernstein, B. E. (2015). Genetic and epigenetic fine mapping of causal autoimmune disease variants. Nature, 518(7539), 337–43.

Gene, P. (2017). Human protein atlas. [Online; accessed 14-August-2017].

Gjoneska, E., Pfenning, A. R., Mathys, H., Quon, G., Kundaje, A., Tsai, L. H., and Kellis, M. (2015). Conserved epigenomic signals in mice and humans reveal immune basis of alzheimer’s disease. Nature, 518(7539), 365–9.

Jaccard, P. (1901). Distribution de la flore alpine dans le bassin des dranses et dans quelques rgions voisines. Bulletin de la Socit Vaudoise des Sciences Naturelles, 37, 241–272.

Ji, Z. (2017). Scrat: Single-cell regulome analysis toolbox. [Online; accessed 3-August-2017].

Langmead, B., Trapnell, C., Pop, M., and Salzberg, S. L. (2009). Ultrafast and memory-efficient alignment of short dna sequences to the human genome. Genome Biol, 10(3), R25.

Li, H., Handsaker, B., Wysoker, A., Fennell, T., Ruan, J., Homer, N., Marth, G., Abecasis, G., Durbin, R., and,. G. P. D. P. S. (2009). The sequence alignment/map format and samtools. Bioinformatics, 25(16), 2078–2079.

Lin, L., Bass, A. J., Lockwood, W. W., Wang, Z., Silvers, A. L., Thomas, D. G., Chang, A. C., Lin, J., Orringer, M. B., Li, W., Glover, T. W., Giordano, T. J., Lam, W. L., Meyerson, M., and Beer, D. G. (2012). Activation of gata binding protein 6 (gata6) sustains oncogenic lineage-survival in esophageal adenocarcinoma. Proceedings of the National Academy of Sciences, 109(11), 4251–4256.

Macaulay, I. C. and Voet, T. (2014). Single cell genomics: advances and future perspectives. PLoS Genet, 10(1), e1004126.

McLean, C. Y., Bristor, D., Hiller, M., Clarke, S. L., Schaar, B. T., Lowe, C. B., Wenger, A. M., and Bejerano, G. (2010). GREAT improves functional interpretation of cis-regulatory regions. Nat Biotechnol, 28(5), 495–501.

Perez, F. and Granger, B. E. (2007). IPython: a system for interactive scientific computing. Computing in Science and Engineering, 9(3), 21–29.

Picard (2017). Picard. [Online; accessed 3-August-2017].

Pliner, H., Packer, J., McFaline-Figueroa, J., Cusanovich, D., Daza, R., Srivatsan, S., Qiu, X., Jackson, D., Minkina, A., Adey, A., Steemers, F., Shendure, J., and Trapnell, C. (2017). Chromatin accessibility dynamics of myogenesis at single cell resolution. bioRxiv.

Quinlan, A. R. and Hall, I. M. (2010). Bedtools: a flexible suite of utilities for comparing genomic features. Bioinformatics, 26(6), 841–842.

Schep, A. N., Wu, B., Buenrostro, J. D., and Greenleaf, W. J. (2017). chromvar: inferring transcription-factor-associated accessibility from single-cell epigenomic data. Nat Meth, advance online publication, –.

Schwartzman, O. and Tanay, A. (2015). Single-cell epigenomics: techniques and emerging applications. Nat Rev Genet, 16(12), 716–26.

Shalek, A. K., Satija, R., Adiconis, X., Gertner, R. S., Gaublomme, J. T., Raychowdhury, R., Schwartz, S., Yosef, N., Malboeuf, C., Lu, D., Trombetta, J. J., Gennert, D., Gnirke, A., Goren, A., Hacohen, N., Levin, J. Z., Park, H., and Regev, A. (2013). Single-cell transcriptomics reveals bimodality in expression and splicing in immune cells. Nature, 498(7453), 236–40.

Shen, H. (2014). Interactive notebooks: Sharing the code. Nature, 515(7525), 151–2.

Thomasset, S. C., Garcea, G., and Berry, D. P. (2008). Oesophageal metastasis from colorectal cancer. Case Reports in Gastroenterology, 2(1), 40–44.

Trimmer, C., Bonuccelli, G., Katiyar, S., Sotgia, F., Pestell, R. G., Lisanti, M. P., and Capozza, F. (2013). Cav1 suppresses tumor growth and metastasis in a murine model of cutaneous scc through modulation of mapk/ap-1 activation. The American Journal of Pathology, 182(3), 992–1004.

Tsompana, M. and Buck, M. J. (2014). Chromatin accessibility: a window into the genome. Epigenetics & Chromatin, 7(1

van der Maaten, L. and Hinton, G. (2008). Visualizing high-dimensional data using t-sne. Journal of Machine Learning Research, 9, 2579–2605.

Zamanighomi, M., Lin, Z., Daley, T., Schep, A., Greenleaf, W. J., and Wong, W. H. (2017). Unsupervised clustering and epigenetic classification of single cells. bioRxiv.

Zhang, Y., Liu, T., Meyer, C. A., Eeckhoute, J., Johnson, D. S., Bernstein, B. E., Nusbaum, C., Myers, R. M., Brown, M., Li, W., and Liu, X. S. (2008). Model-based analysis of chip-seq (macs). Genome Biology, 9(9), R137.

